# Discovery of rhodomyrtone as a broad-spectrum antiviral inhibitor with anti-SARS-CoV-2 activity

**DOI:** 10.1101/2020.11.14.382770

**Authors:** Wei Tang, Jing Lu, Qiao-Yun Song, Man-Mei Li, Li-Feng Chen, Li-Jun Hu, Shao-Min Yang, Dong-Mei Zhang, Ying Wang, Yao-Lan Li, Wen-Cai Ye

## Abstract

The outbreak of new viruses, such as serve acute respiratory syndrome coronavirus 2 (SARS-CoV-2), as well as the emerging of drug-resistance viruses highlight the urgent need for the development of broad-spectrum antiviral drugs. Herein, we report the discovery of a plant-derived small molecule, 6,8-dihydroxy-9-isobutyl-2,2,4,4-tetramethyl-7-(3-methylbutanoyl)-4,9-dihydro-1*H*-xanthene-1,3(2*H*)-dione (rhodomyrtone, RDT), which exhibited potent broad-spectrum antiviral activities against several RNA and DNA viruses, including SARS-CoV-2, respiratory syncytial virus (RSV), herpes simplex virus type 1 (HSV-1), herpes simplex virus type 2 (HSV-2), varicella-zoster virus (VZV), human cytomegalovirus (HCMV), and Kaposi’s sarcoma-associated herpesvirus (KSHV). RDT can significantly suppress viral gene expression and show the low possibility to elicit drug-resistant variants. Mechanistic study implied that RDT inhibited viral infection by disturbing the cellular factors that essential for viral gene expression. Our results suggested that RDT might be a promising lead compound for the development of broad-spectrum antiviral drugs.

---

Most of the current drugs are used to treat virus infections by directly targeting viral factors, particularly the enzymes (e.g., protease or DNA/RNA polymerase)^1,2^. In the past decades, these direct-acting drugs have achieved great success to fight virus-associated diseases. However, a variety of issues, such as drug resistance and adverse effects, have been developed^2^. Moreover, a direct-acting antiviral drug is often effective in inhibiting the viruses of the same subfamily, it is incapable when the patients are infected with multiple viruses or undiagnosed viruses. In contrast, the broad-spectrum antiviral inhibitors would be a great significance in these cases. In addition, with the rapid emergence of new viruses in recent years, the development of broad-spectrum antiviral drugs is urgently needed. Recently, a novel severe acute respiratory syndrome coronavirus 2 (SARS-CoV-2) has emerged and is rapidly spreading around the world, resulting in over 40.9 million infections and 1,125,474 deaths until now. Currently, the therapeutics for SARS-CoV-2 are still limited, and most of the drug candidates against SARS-CoV-2 target the viral factors^3^. With the rapid mutation in the genome of SARS-CoV-2^4,5^, the direct-acting antiviral drugs would increase the risk of resistance.

Search for the broad-spectrum antiviral drugs targeting host cell factors essential for the reproduction of virions is becoming an alternative approach. Also, broad-spectrum antiviral drugs minimize the risk of drug resistance. As an ongoing research interest of our group in the past decades, we have constructed an in-house compound library composing of structural diversity natural products obtained from approximately 100 species of medicinal plants. After systematic bioassay screening, a series of compounds were found to exhibit potent antiviral activities^6–10^. In this paper, we report a plant-derived small molecule, 6,8-dihydroxy-9-isobutyl-2,2,4,4-tetramethyl-7-(3-methylbutanoyl)-4,9-dihydro-1*H*-xanthene-1,3(2*H*)-dione (rhodomyrtone, **Fig. 1a**), with broad-spectrum antiviral activities^11,12^.

**Fig. 1.**
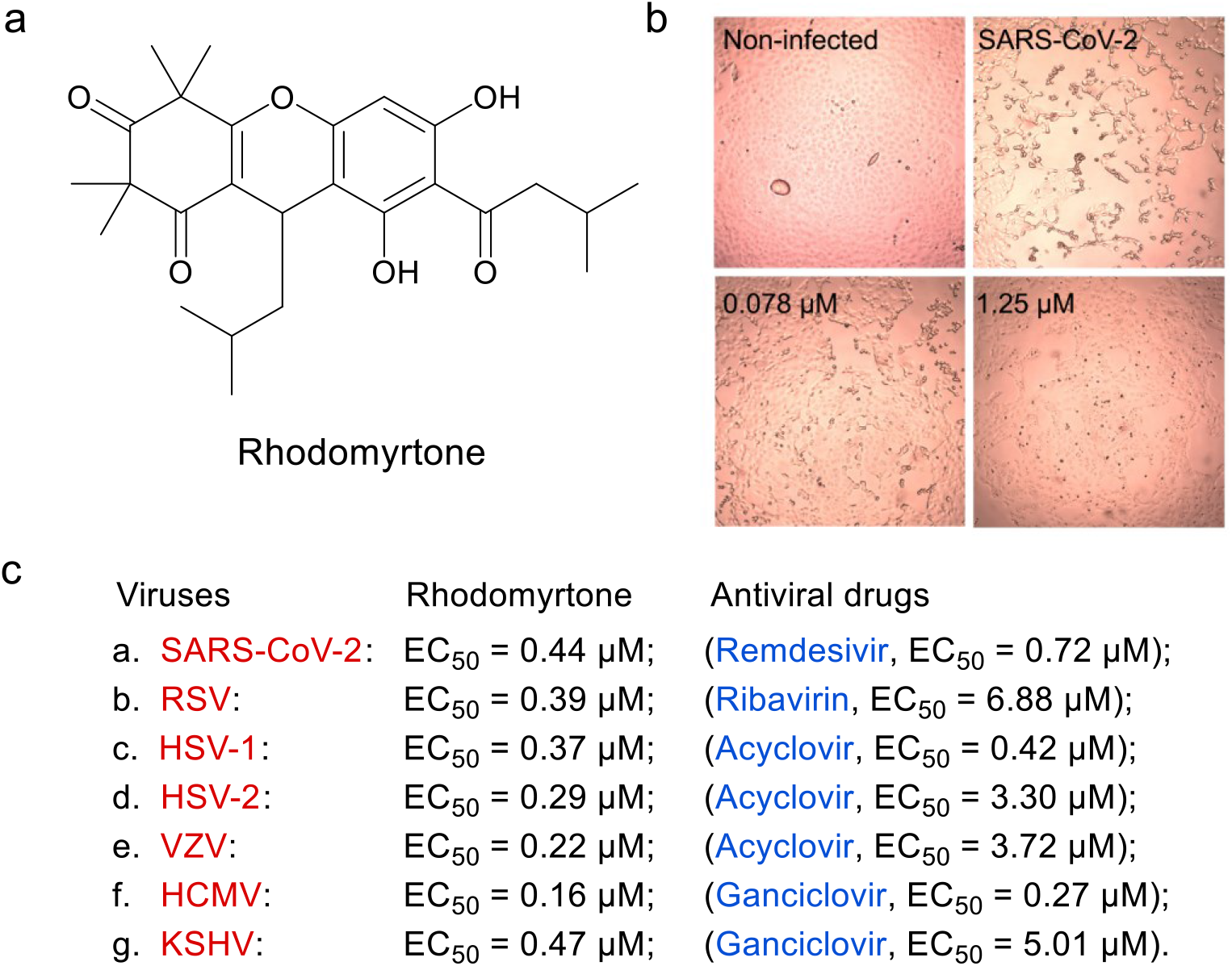
Antiviral activities of rhodomyrtone (RDT) against different viruses. **a**, Chemical structure of RDT. **b**, Inhibition of SARS-CoV-2-induced cytopathic effect (CPE) by RDT. Vero E6 cells were infected with SARS-CoV-2 and treated with RDT at different concentrations. At 48 h after the viral infection, the CPE in the cells were observed and photographed under a microscope. **c**, Inhibitory effect of RDT or FDA approved antiviral drugs against different viruses.

Rhodomyrtone (RDT), a phloroglucinol dimer originally isolated from traditional Chinese medicine “Tao-Jin-Niang” (*Rhodomyrtus tomentosa* (Ait.) Hassk.), was previously reported to be a Gram-positive bacteria inhibitor that could significantly inhibit the growth of *Staphylococcus aureus* by targeting the bacterial farR protein^13^. However, the antiviral activities of RDT have not been explored. In the present study, we demonstrated for the first time that RDT potently inhibited several DNA and RNA viruses (**Fig. 1b–c**), including severe acute respiratory syndrome coronavirus 2 (SARS-CoV-2), respiratory syncytial virus (RSV), herpes simplex virus type 1 (HSV-1), herpes simplex virus type 2 (HSV-2), varicella-zoster virus (VZV), human cytomegalovirus (HCMV), and Kaposi’s sarcoma-associated herpesvirus (KSHV). *In vitro*, the antiviral activities of RDT are stronger than those of several FDA-approved nucleoside analogue antiviral drugs, such as remdesivir, ribavirin, acyclovir (ACV), and ganciclovir (GCV). Importantly, RDT potently suppressed SARS-CoV-2 infection in Vero E6 cells at the concentrations without exhibiting substantial cytotoxicity. The EC_50_ of RDT against SARS-CoV-2 is 0.44 μM, which is lower than that of remdesivir.

Subsequently, the antiviral mechanism of RDT against HSV-1 was investigated. We found that HSV-1/Blue virus, a TK mutant variant, showed strong resistance to ACV but sensitive to RDT. The virus-induced plaques were significantly inhibited by the treatment of RDT at the concentrations from 0.5 to 2 μM, however, the plaque formation was not affected by ACV even at the concentration of 20 μM (**Fig. 2a**). In time-of-addition assay, the reproduction of HSV-1 was almost completely suppressed by 1 μM of RDT as well as 2 μM of ACV at 0, 2, 4, 6 h post-infection. The inhibitory effect of ACV on HSV-1 was diminished at 8 h post-infection, however, the virus in the cells was still potently inhibited by RDT until 14 h after viral infection (**Fig. 2b**). These results indicated that the action mechanism of RDT is distinct from that of ACV.

**Fig. 2.**
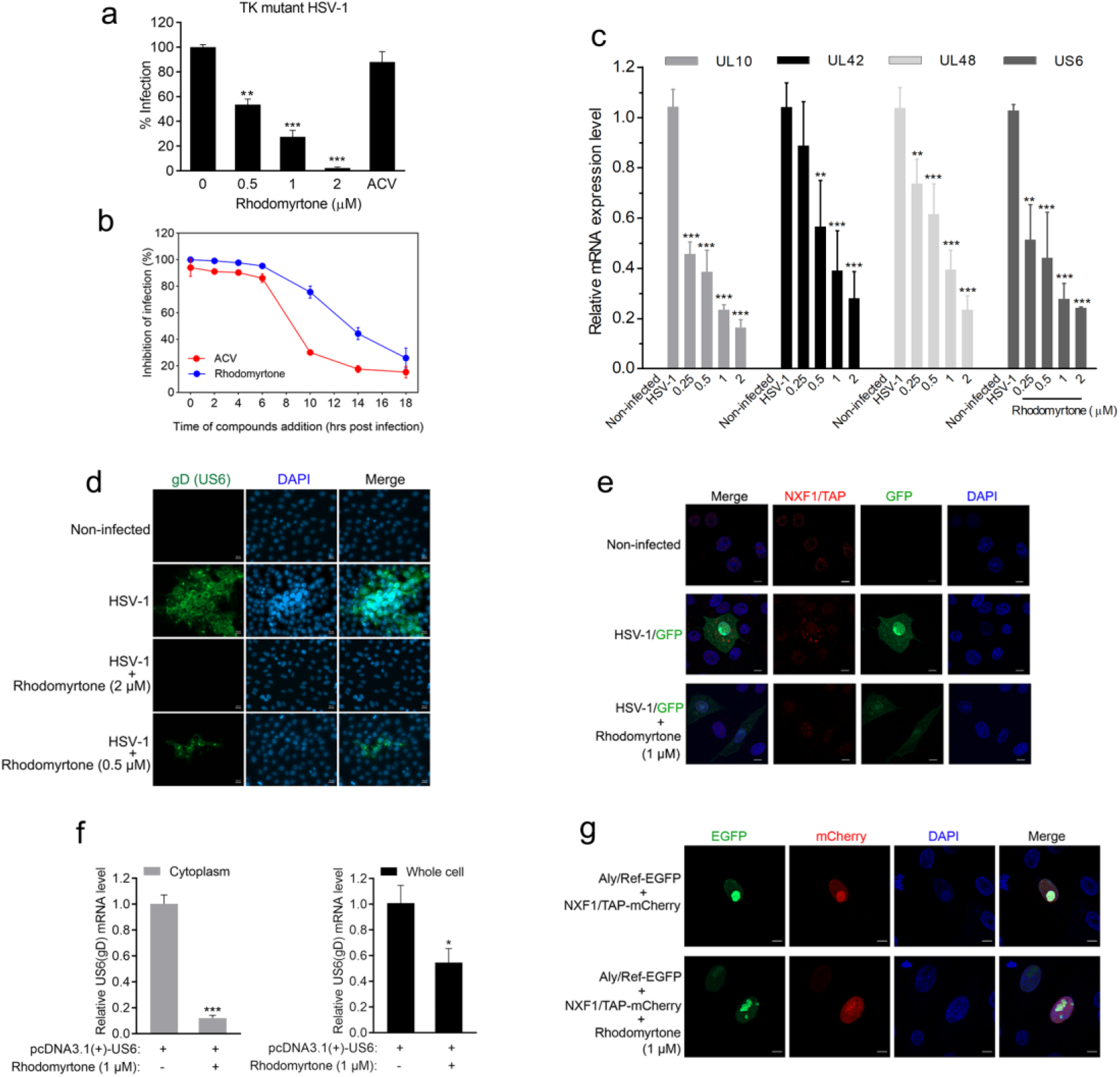
Antiviral mechanism of RDT against HSV-1. **a**, Inhibition of ACV-resistant strain (HSV-1/Blue) infection by ACV (20 μM) or RDT at different concentrations. **b**, Inhibition of HSV-1/F infection by the treatment of RDT (1 μM) or ACV (2 μM) to the Vero cells at different time points after viral infection. **c**, **d**, Inhibitory effect of RDT on viral gene expression. The transcripts levels of viral genes were quantified by RT-PCR (**c**). The gD proteins of HSV-1/F in Vero cells were incubated with anti-gD monoclonal antibody, followed by staining with Alexa Fluor 488 secondary antibody (**d**). **e**, Vero cells were infected with HSV-1/Patton/GFP with or without treatment of RDT. The cellular NXF1/TAP was incubated with a specific anti-NXF1/TAP antibody and then stained with Alexa Fluor 594 secondary antibody. Scale bar, 10 μm. **f**, The mRNA level of viral US6 (gD) in the nucleus or cytoplasm after treatment of RDT. HEK293T cells were transfected with pcDNA3.1(+)-gD plasmids with or without the treatment of RDT. **g**, Plasmids of pEGFP-Aly/Ref and pmCherry-NXF1/TAP were cotransfected to HEK293T cells, followed by the treatment of RDT at 1 μM. The co-localization between Aly/Ref and NXF1/TAP was observed and photographed by using a confocal microscope. Scale bar, 10 μm.

The results of immunofluorescence and RT-PCR assay demonstrated that RDT potently suppressed the viral gene expression in a concentration-dependent manner (**Fig. 2c-d**). Interestingly, the transcription level of US6 (gD) that inserted in a pcDNA3.1(+) plasmid was decreased after the treatment of RDT in the cells, particularly significantly decreased in the cytoplasm (**Fig. 2e-f**). Next, we observed that the distribution of NXF1/TAP, a cellular mRNA export receptor, in the cytoplasm was inhibited by RDT treatment, suggesting an inhibitory effect of RDT on mRNA export (**Fig. 2g**). To determine whether the mRNA export pathway was affected by RDT, we inserted Aly/Ref, a cellular mRNA export adaptor, and NXF1/TAP into pEGFP-N3 and pmCherry-N1 plasmids, respectively. The results in confocal assay demonstrated that NXF1/TAP-mCherry was co-localized with Aly/Ref-EGFP in the cells, however, their interaction was affected by the treatment of RDT (**Fig. 2h**). Together, these results implied that RDT is a broad-spectrum antiviral inhibitor targeting the cellular factors involved in viral gene expression. Moreover, a resistant variant was not be generated despite culturing the virus in the presence of RDT over two months, providing evidence that RDT is not a direct-acting antiviral inhibitor.

In summary, RDT is a promising inhibitor against SARS-CoV-2 with broad-spectrum antiviral activities. It inhibits viral infection by affecting the distribution of cellular factors essential for viral gene expression. Further research including *in vivo* antiviral activity and potential target of RDT against different viruses are remaining to be elucidated.

## Materials and Methods

### Cells, viruses, and compounds

African green monkey kidney epithelial cells (Vero, ATCC) and HEK293 T cells (ATCC) were cultured in DMEM containing 10% fetal bovine serum (FBS, Gibco Invitrogen) at 37 °C, 5% CO_2_. The viruses used in this study were SARS-CoV-2/Guangdong/20SF014, RSV/A2 HSV-1/F, HSV-1/Patton/GFP, HSV-2/MS/GFP, HCMV/Towne/GFP, KSHV/BAC16/GFP, and VZV/ROka. All the viruses were separated into aliquots and stored at −70 °C. Rhodomyrtone (RDT) was isolated from the leaves of *Rhodomyrtus tomentosa* (Ait.) Hassk. and identified as 6,8-dihydroxy-9-isobutyl-2,2,4,4-tetramethyl-7-(3-methylbutanoyl)-4,9-dihydro-1H-xanthene-1,3(2H)-dione by nuclear magnetic resonance spectroscopy (NMR), and the purity of which was over 99% analyzed by high-performance liquid chromatography (HPLC). All tested compounds, including RDT, acyclovir (ACV), ganciclovir (GCV), and ribavirin were dissolved in DMSO.

### Plasmids

The full lengths of cellular Aly/Ref and NXF1/TAP genes were cloned into pEGFP-N3 and pmCherry-N1 vectors, respectively. The C-terminus of Aly/Ref was fused with the N-terminus of EGFP fragment in pEGFP-N3, and the C-terminus of NXF1/TAP was fused with the N-terminus of mCherry fragment in pmCherry-N1. The fused proteins Aly/Ref-EGFP and NXF1/TAP-mCherry were transiently expressed in HEK293T cells by transfection with lipofectamine 2000. The viral US6 (gD) gene was cloned into a pcDNA3.1(+) vector.

### Time of addition assay

Vero cells were seeded in 96-well culture plates at the density of 1×10^4^ cells per well and incubated overnight. Following the incubation, the cells were infected with HSV-1/F at a multiplicity of infection (MOI) of 0.2 and treated with ACV (2 μM) or RDT (1 μM) at indicated time points after viral infection. At 24 h post-infection, the virus suspensions were harvested by frozen-thawed at −70 °C and centrifuged at 2000×g for 10 min. Viral titers in the prepared supernatants were determined by plaque reduction assay^7^.

### Immunofluorescence assay

Vero cells were seeded in 96-well culture plates and incubated overnight. The cells were then infected with HSV-1/F (MOI=0.2) and treated with RDT at different concentrations. At 24 h after the viral infection, the cells were fixed with 4% formaldehyde in PBS for 30 min and washed twice with PBS, followed by the permeabilized treatment with 0.1% Triton X-100 for 10 min. The cells were then washed with PBS twice, blocked by 5% BSA and incubated with mouse anti-HSV-1 gD monoclonal antibody (1:500, Santa Cruze Biotechnology) for 2 h, and stained with Alexa Fluor 488 anti-mouse secondary antibody (1:1000, Thermo Fisher Scientific) for 1 h at 4 °C. After washing with PBS twice, the cells were performed on a fluorescence microscope.

### RT-PCR

Vero cells in 12-well culture plates were infected with HSV-1/F at MOI = 0.5 or transfected with pcDNA3.1(+)-US6(gD) plasmid, and then treated with RDT at various concentrations. After infection at 4 °C for 1 h, the cells were washed with PBS twice to remove the unbound virus and supplemented with the culture medium containing different concentrations of RDT. At 9 h post-infection or 12 h after transfection, total RNA in the cells were harvested and purified by using ion-exchange minicolumns (RNA fast 200 kits, Fastagen), following the manufacturer’s instructions. Total RNA (1 μg) was used as a template for cDNA synthesis by RT-PCR. The cDNA templates were then used for Sybr green real-time PCR to determine the transcription level of viral genes with a Light Cycler 480 system (Roche).

### Confocal assay

Vero cells in confocal dishes were infected with HSV-1/Patton/GFP (MOI=0.2), and then treated with RDT at 1 μM. At 12 post-infection, the cells were fixed by 4% paraformaldehyde in PBS and permeabilized with 0.05% Triton X-100 for 15 min. After washing 3 times with PBS, the cells were incubated with rabbit anti-NXF1/TAP primary antibody (1:500, Cell Signaling Technology) and Alexa Fluor 594 anti-rabbit secondary antibody (1:500, Thermo Fisher Scientific), followed by staining the nucleus with DAPI. For determining the co-localization between Aly/Ref and NXF1/TAP, HEK293T cells were cotransfected with pEGFP-Aly/Ref and pmCherry-NXF1 plasmids with lipofectamine 2000 (Thermo Fisher Scientific). Following the transfection, the cells were treated with the RDT at 1 μM. At 12 h after transfection, the cells stained with DAPI and performed on a confocal laser microscope (Zeiss LSM800).

### Statistical analysis

Data are presented as the mean ± SD unless indicated otherwise. Statistical analyses were conducted using Prism, version 8.0. Student’s T-test (two-tailed assuming equal variances) was used to compare groups. A value of P < 0.05 was considered statistically significant.

## Acknowledgments

This research has been supported by grants from the Key-Area Research & Development Program of Guangdong Province – Special Program for COVID-19 (No. 202020012620800001), the Local Innovative & Research Teams Project of Guangdong Pearl River Talents Program (No. 2017BT01Y036), and the Key-Area Research & Development Program of Guangdong Province (No. 2020B1111110004).

